# A parvocellular-magnocellular functional gradient in human visual cortex

**DOI:** 10.1101/2022.02.08.479509

**Authors:** Maddi Ibarbia-Garate, Pedro M. Paz-Alonso

## Abstract

The magnocellular and parvocellular systems are major visual recognition pathways, with distinct histological and physiological properties. Despite their critical role, there is limited evidence on the specific contributions these visual pathways make to visual recognition in general and to word reading in particular. Using a multimodal functional MRI approach and individual-subject analyses, here we investigate the involvement of visual cortex regions in the recognition of magnocellular- and parvocellular-biased words and images. Our results reveal a functional gradient in the activation profile of both left and right visual cortex: posterior regions are more strongly recruited for processing parvocellular-biased, while anterior regions are more involved in processing magnocellular-biased stimuli. Furthermore, functional connectivity analyses show clustering in the strength of functional coupling among visual cortex regions as a function of the distance between regions, with greater coupling within and less coupling across posterior and anterior regions. Finally, we found minimal differences in lateralization for word and image recognition in these visual cortical regions. These results were replicated in a retest session with a subset of participants. Our findings underscore a functional division of labor in the visual cortex as a function determined by a parvocellular or magnocellular bias in properties of the stimuli and further reveal that in this context the visual cortex is not particularly biased towards words or images.

## 1. Introduction

Object recognition and reading are activities we perform on a daily basis. These operations are complex and engage a cascade of neural processes that need to be orchestrated in an organized, sequential manner, including in the initial step: visual recognition. Visual input is analyzed by a hierarchy of regions situated along the dorsal and ventral visual streams wherein the magnocellular and parvocellular systems play critical and functionally distinct roles. The magnocellular dorsal pathway is achromatic, sensitive to higher contrast stimuli with lower spatial and higher temporal frequencies, and exhibits transient responses (e.g., Ungerleider and Haxby, 1994). By contrast, the parvocellular ventral pathway is color sensitive, responds to lower contrast stimuli with higher spatial and lower temporal frequencies, and exhibits sustained responses (e.g., Merigan, Katz, & Maunsell, 1991).

Most models of visual and object recognition concur that visual analysis starts with the parallel extraction of different elementary visual attributes at different spatial frequencies (e.g., Bullier, 2001; Hegdé, 2008; Peyrin et al., 2010; Schyns & Oliva, 1994), and that visual recognition encompasses feed-forward bottom-up processes (e.g., Grossberg, 1982; Kosslyn, 1994), as well as facilitative top-down processes (e.g., Bar, 2003; Barceló et al., 2000; Bullier, 2001; Miyashita et al., 2000). Although it has been suggested that similar related processes occur in the visual cortex during reading (Rauschecker et al., 2011), there is still limited evidence on the specific contributions the magnocellular and parvocellular systems make to reading and object recognition.

Previous evidence indicates that the magnocellular dorsal pathway is the dominant visual pathway for text perception (Chase et al., 2003) and reading (Conlon et al., 2004). This work suggests that neural structures within this pathway may play a role in both motion processing and visual language processing. Further, word recognition performance and magnocellular-related abilities have been positively correlated (Ben Shachar et al., 2007, Laycock et al., 2009, Oludade et al., 2013, Joo et al., 2017) and training on magnocellular- related tasks (e.g., watching moving versus stationary dots) can improve visual word recognition (Chouake et al., 2012). In addition, research on developmental dyslexia – a neurobiological reading disorder that can occur despite normal intelligence, adequate education, and lack of obvious sensory or neurological damage (American Psychiatric Association, 2013; World Health Organization, 2008) – suggests some involvement of the magnocellular pathway (e.g., Livingstone et al., 1991; Stein & Walsh, 1997; Stein, 2001; Franceschini et al., 2012, Gori et al., 2015). Specifically, some children with reading disabilities appear to suffer from weakened or abnormal magnocellular input to the dorsal visual pathway, causing dysfunction of the main frontoparietal attentional network (Livingstone et al., 1991, Stein et al., 1997). There is less evidence that the parvocellular ventral pathway plays a role in reading (but see for instance Farrag et al., 2002). But this may be partly due to the fact that previous studies exclusively focused on the magnocellular pathway without including specific manipulations that could have revealed involvement of the parvocellular pathway in reading processes in particular, and visual recognition in general.

Within the visual cortex, the more posterior regions (i.e., V1, V2, and V3) receive inputs from both the magnocellular and parvocellular pathways (Rauschecker et al., 2011). In contrast, hv4 is thought to be a color selective region, and therefore more prone to respond to parvocellular-related processes (Zeki 1980; Zeki 1983, Vaina 1994; Bartels and Zeki 2000; Brewer et al. 2005). Lateral occipital (LO) regions are considered part of the object selective lateral occipital complex (Malach et al., 1995; Grill-Spector et al., 2000) and the LO1 region tends to show orientation-selective responses to simple grating stimuli (Larsson et al., 2006). Finally, temporal occipital (TO) regions (TO1 and TO2) fall within the motion-selective cortex, also called MT+/V5 (Dumoulin et al., 2000, Amano et al., 2009), and are involved in motion perception (Tootell et al., 1995, Rees et al., 2000, Grill-Spector et al., 2004). Due to their sensitivity to orientation and motion, the LO1 and TO regions are thought to be especially relevant for processing magnocellular stimuli (Maunsell et al., 1990). Previous research on low versus high spatial frequency processing has shown that spatial frequencies are mapped in a retinotopic manner (Fox et al., 1987, Engel, Glover, & Wandell, 1997, Sasaki et al., 2001) and suggest that low spatial frequencies are processed in the anterior part of V1, whereas high spatial frequencies are processed in the posterior and ventral parts of V2, V3, V4 (Mussel et al., 2013).

The present fMRI study was aimed at characterizing the functional contributions of the magnocellular and parvocellular visual pathways to reading, in particular, as well as to visual recognition of object images, more generally. Based on previous evidence, we hypothesized that 1) peripheral or more anterior visual cortex regions would be more strongly recruited during the processing of magnocellular-biased stimuli (Maunsell et al., 1990, Tootell et al., 1995, Rees et al., 2000, Grill-Spector et al., 2004); 2) since evidence for involvement of early visual cortex regions in processing parvocellular- or magnocellular- biased stimuli is limited, and to some extent mixed, we wanted to further explore whether these regions would be differentially recruited for processing magnocellular-biased or parvocellular-biased words and images; 3) in line with hypotheses 1 and 2, we expected tighter functional connectivity within than between posterior and anterior visual cortex regions; finally, 4) we predicted laterality effects for visual recognition of letter strings in left (versus right) peripheral or anterior visual cortex regions typically associated with magnocellular operations (Ray et al., 2005) in line with left hemisphere involvement in language processing (Lau et al., 2008).

## 2. Materials and Methods

### 2.1. Participants

The study sample consisted of 34 participants (19 females and 15 males, mean age 25.37 ± 4.41 years). Fourteen of them (8 females and 6 males, mean age 25.10 ± 4.58 years) also participated in an identical second session 7–10 days later (i.e., retest session), to examine data reproducibility (see Figure 3, 4 and 5 supplement 2). Three additional participants were excluded from further analyses due to excessive head motion during scanning (see “*MRI data acquisition and analysis*” section below). Also, one additional participant was excluded due to low behavioral performance on the fMRI task (less than 60% accuracy).

All participants were right-handed and had normal or corrected-to-normal vision. No participant had any history of major medical, neurological, or psychiatric disorders. The study protocol was approved by the Ethics Committee of the Basque Center on Cognition, Brain and Language (BCBL) and was carried out in accordance with the Code of Ethics of the World Medical Association (Declaration of Helsinki) for experiments involving humans. Prior to their inclusion in the study, all subjects provided informed written consent. Participants received monetary compensation for their participation.

### 2.2. Stimuli

A total of 180 line drawings of words and images were used as stimuli. The uniform foreground of these drawings allows for precise control of luminance and chromatic properties. Half were magnocellular-biased, that is achromatic with low-luminance contrast; the other half were parvocellular-biased, that is, chromatically defined and isoluminant (red-green).

It is known that tuning of the magnocellular and parvocellular visual pathways differs across individuals, so we used standard procedures to adjust luminance and chromatic settings before MRI scanning (Cheng et al., 2004; Kveraga et al., 2007; see also Skottun, 2013, for cautions on using isoluminant stimuli). The standard technique for establishing the luminance threshold for magnocellular-biased achromatic stimuli involves a multiple staircase procedure, during which subjects are required to answer whether or not they can identify image and word stimuli. Once the luminance threshold has been established, the appropriate luminance (∼3.5% Weber contrast) value is defined for the gray-scale line drawings to be used in the low-luminance-contrast, magnocellular-biased, condition. We used a common luminance value across all participants for the magnocellular-biased stimuli; we had shown that these stimuli met an 80% accuracy criterion in a normative study (N = 15) made prior to the current experiment. In that normative study, we used 40 words and 40 images (i.e., line drawings corresponding to each of these words), in a counterbalanced fashion, to determine the appropriate luminance contrast for participants to reach 80% accuracy on the achromatic magnocellular-biased stimuli. The criterion of minimum 80% of accuracy was used to guarantee enough correct responses to magnocellular-biased stimuli, since magnocellular-biased trials are, in general, more difficult to process than parvocellular-biased ones.

For the parvocellular-biased, chromatically defined stimuli, the isoluminance point was identified for each subject individually using heterochromatic flicker photometry as reported in previous studies (Kveraga et al., 2007). This procedure identifies the isoluminance point for two colors by having a stimulus rapidly alternate between them. The color values at which the stimuli appears to stop flickering indicates the narrow isoluminance interval of the steady point. Each participant was required to indicate the steady point for line drawings of images displayed in pure red and pure green before undergoing MRI scanning (*M* = 136.5 ± 26.16 in the initial session; M = 122 ± 18.50 in the retest session).

All of the images and word stimuli were designed in both a magnocellular-biased and a parvocellular-biased form and each word had a corresponding image, so that all stimuli were properly counterbalanced across subjects in terms of Pathway (i.e., magnocellular, parvocelullar) and Stimuli (i.e., words, images) (see Figure 1). All the selected items could be easily identified as non-manmade (i.e., *apple*) or manmade (i.e., *robot*); 50% of items were “non-manmade”, and the remaining 50% were “manmade”.

**Figure 1.**
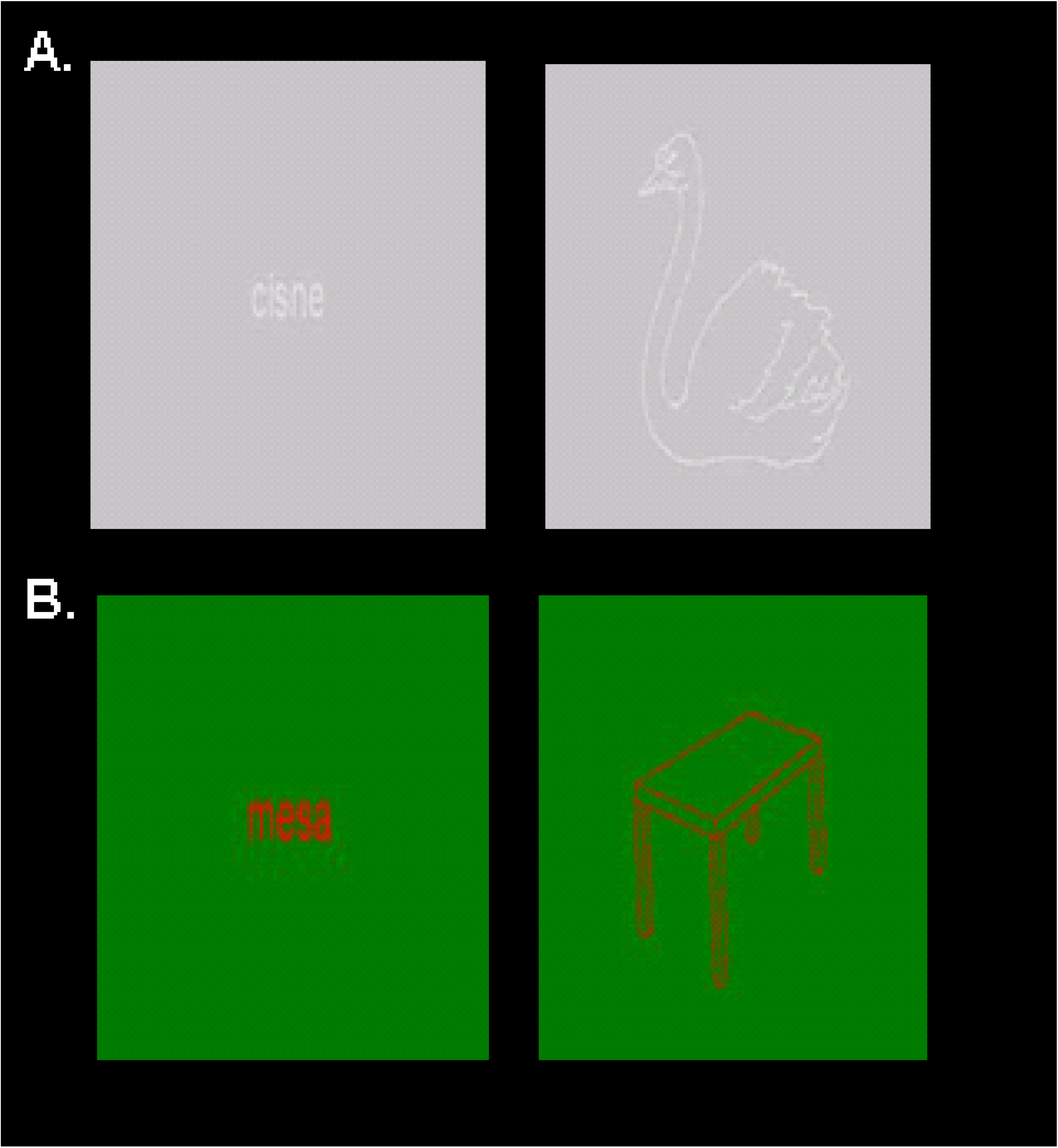
Experimental stimuli. We used a hybrid event/block design where participants made manmade (i.e., artificial) or non-manmade (i.e., natural) judgments on words and images that were A) magnocellular biased, with low luminance and contrast, and achromatic; or B) parvocellular biased; isoluminant (red-green) and chromatically defined. All stimuli were counterbalanced across conditions and subjects.

### 2.3. Experimental task and procedure

Functional and structural data was collected in the MRI scanner (see *MRI data acquisition* section below). The functional MRI task conformed to a 2 (Pathway: magnocellular, parvocellular) X 2 (Stimuli: words, images) visual recognition experimental design, which was carried out in a single functional run that lasted 15 min. During the task, participants were asked to perform a semantic task, in which they indicated natural (i.e., non man-made) or artificial (i.e., man-made) judgements via key presses on a two-button fiber-optic box. Six stimuli were presented in each activation block of 20.3 s duration. Activation blocks were alternated with rest-fixation periods of 16 s to allow the hemodynamic response function (HRF) to return to baseline. A total of 90 words and 90 images were presented across activation blocks, yielding 5 activation blocks for each of the 4 experimental conditions, plus 2 additional blocks of baseline conditions that were not included in subsequent analyses. The stimuli were presented at 100 Hz using a Barco projector (model F35); the refresh rate was confirmed using The Black Box ToolKit v2™.

For behavioral analysis, we conducted two separate 2 (Pathway: magnocellular, parvocellular) X 2 (Stimuli: words, images) repeated measures analyses of variance (ANOVA) with accuracy and reaction times as dependent measures. Only blocks with at least 60% correct responses were included in the behavioral and fMRI analyses (total number of excluded blocks from the final sample [N = 34] = 69; mean excluded blocks per subject = 0.06). We used the mean ± 2.5SD criterion for outliers in all measures in this study.

We applied multiple comparison corrections to all behavioral as well as functional MRI results. We also computed effect sizes and Bayes factor values (BF10 > 3 suggests substantial evidence for a difference between pairs; BF10 < 0.3 suggests substantial evidence for a null effect, see Jeffreys, 1961).

### 2.4. MRI data acquisition and analysis

Whole-brain MRI data acquisition was conducted on a 3-T Siemens PRISMA Fit whole-body MRI scanner (Siemens Medical Solutions) using a 64-channel whole-head coil. Functional images were acquired in a single gradient-echo echo-planar multiband pulse sequence with the following acquisition parameters: time-to-repetition (TR) = 1000 ms; time-to-echo (TE) = 35 ms; MB acceleration factor = 5; 65 axial slices with a 2.4 mm^3^ voxel resolution; no inter-slice gap; flip angle = 56°; field of view (FoV) = 210 mm; 865 volumes. High-resolution MPRAGE T1-weighted structural images were also collected for each participant with the following parameters: TR = 2530 ms; TE = 2.36 ms; flip angle = 7°; FoV = 256 mm; voxel resolution = 1 mm^3^; 176 slices.

For *structural analysis* of the T1-weighted images, we used the Freesurfer 6.0 pipeline (Fischl et al., 2004) to perform volumetric gray and white matter segmentation, providing several automated cortical parcellations for use in subsequent analyses. After obtaining the gray-matter segmentation with Freesurfer, we ran Benson’s atlas (Benson et al., 2014) to obtain all of the regions of the visual cortex included in this atlas at the individual-subject level.

For *functional data preprocessing and analysis,* we used SPM12 (Wellcome Center for Human Imaging, London) preprocessing routines and analysis methods. Images were corrected for differences in slice acquisition timing across every functional scan and then realigned for motion correction. Afterwards, each subject’s functional volumes were smoothed using a 2-mm full-width half-maximum (FWHM) Gaussian kernel. Motion parameters were extracted from the realignment step to inform a volume repair procedure (ArtRepair; Stanford Psychiatric Neuroimaging Laboratory) that identified bad volumes on the basis of scan-to-scan movement (>0.5 mm) and signal fluctuations in global intensity (>1.3%) and corrected bad volumes via interpolation from the nearest non-repaired scans. One participant with more than 20% to-be-corrected outlier volumes in the functional run was excluded. Two participants were also excluded due to motion during structural T1- weighted images acquisition preventing accurate structural-functional overlap after corregistration. For the final sample of participants, the average percentage of repaired volumes was 2% (SD = 3%). After volume repair, high-resolution anatomical T1 images and functional volumes were coregistered and resliced from the original 2.4 x 2.4 x 2.4 mm functional voxel dimensions to 1 x 1 x 1 mm voxels in anatomical T1 space. Finally, time series were temporally filtered to eliminate contamination from slow frequency drift (high-pass filter: 128 s). Once all functional images were in the same space as the individual anatomical images, we ran Benson’s Neuropythy tool (Benson et al., 2014) to segment the main visual cortex regions at the individual-subject level.

Statistical analyses were performed on individual-subject data using the general linear model (GLM). A series of impulses convolved with a canonical hemodynamic response function (HRF) were used to model the fMRI time series data. The four main experimental conditions in our design (i.e., magnocellular-biased words, magnocellular-biased images, parvocellular-biased words, and parvocellular-biased images) were modeled from the onset of the presentation of the first stimulus within each block until the end of the presentation of the last experimental stimulus within the block, resulting in 20.3 s epochs. These functions were used as covariates in the GLM. The motion parameters for translation (i.e., x, y, z) and rotation (i.e., yaw, pitch, roll) were used as covariates of non-interest in the GLM. SPM12 FAST was used for temporal autocorrelation modeling in this GLM due to its optimal performance in terms of removing residual autocorrelated noise in first-level analyses (Olszowy et al., 2019), especially relevant for the present study since data was processed in individual-subject space. The least-squares parameter estimates of the height of the best-fitting canonical HRF for each condition were used in pairwise contrasts. Importantly, blocks with less than 60% correct responses were modeled separately and not considered in the main analyses (the total number of excluded blocks across the final sample for the initial session [N = 34] = 69; mean excluded blocks per subject for the initial session = 0.06; total amount of excluded blocks across the final sample for the retest session [N = 14] = 66; mean excluded blocks per subject for the retest session = 0.010). As previously indicated, an additional participant with more than 10 blocks with less than 60% correct responses was excluded from the final sample.

#### Individual ROI Analysis

ROI analysis was performed with the MARSBAR toolbox for use with SPM12. Given that this study focused on the involvement of visual cortex in visual recognition of letter strings, a total of six left-lateralized visual regions of interest were examined. These regions, extracted individually based on the use of Benson’s atlas with Freesurfer (Benson et al., 2014), were V1, V2, V3, hV4, lateral occipital (LO1), and temporal occipital (TO1). The corresponding right-lateralized regions were also extracted to examine whether they would follow the same functional patterns as the left hemisphere (see Figure 3, supplement 1). Then, in line with our main experimental design, and in order to avoid problems associated with greater signal in more posterior compared to more anterior areas, we extracted parameter estimates (i.e., scaled % signal change values; Mazaika, 2009) and signal t-values for specific contrasts related to the factor Pathway (i.e., Magnocellular > Parvocellular Words; Magnocellular > Parvocellular Images; Parvocellular > Magnocellular Words; Parvocellular > Magnocellular Images) for each single region at the individual-subject level. Finally, 6 (ROI: V1, V2, V3, hV4, LO1, and TO1) X 2 (Stimuli: words, images) repeated measures ANOVAs were performed. Results were corrected using the false discovery rate (FDR) correction for multiple comparisons.

#### Probabilistic Maps

Four surface-based probabilistic maps were calculated using Analysis of Functional Images (AFNI) software (Cox et al., 1996, Cox et al., 1997, Saad et al., 2004) to examine the differential contribution of voxels within visual cortex ROIs (i.e., V1, V2, V3, hv4, LO1, TO1). First, the activations inside the six visual cortex ROIs were binarized for the two contrasts by using a threshold common to all the subjects (i.e., for every subject, all voxels that had a positive t-values were scored 1 and used for the magnocellular-biased maps with the rest zeroed, whereas all voxels that had a negative t-values were scored 1 and used for the parvocellular-biased maps with the rest zeroed). Thus, the binarization step initially yielded four different binarized maps: Magnocellular > Parvocellular Words; Magnocellular > Parvocellular Images; Parvocellular > Magnocellular Words; and, Parvocellular > Magnocellular Images. Due to the lack of significant interactions involving the factor Stimuli in ROI analyses, we also obtained probabilistic maps for Magnocellular > Parvocellular and Parvocellular > Magnocellular contrasts across stimuli conditions. As all ROI and related functional data was in individual-subject space, the binarized maps and the ROIs were normalized to MNI space using the Advanced Normalization Toolbox (ANTs) to obtain these probabilistic maps. After normalization to standard space, corresponding ROIs from every subject were superimposed on the four maps. For each ROI, a probability map was generated by dividing, at each particular voxel, the number of times that voxel belonged to that ROI by the number of subjects included for that ROI. A voxel would have a 100% value if all subjects had this voxel activated for the indicated contrast. A voxel would have a 0% value if none of the subjects had this voxel activated for the indicated contrast.

#### Functional Connectivity Analysis

Functional connectivity analyses were conducted via the beta-series correlation method (Rissman et al., 2004), implemented in SPM12 with custom Matlab scripts. The canonical HRF in SPM was fit to each trial from each experimental condition and the resulting parameter estimates (i.e., beta values) were sorted according to the study conditions to produce a condition-specific beta series for each voxel. Pairwise functional connectivity analysis between each pair of visual cortex ROIs (i.e., V1, V2, V3, hV4, LO1, TO1) were conducted at the individual-subject level. This was done by applying an arc hyperbolic tangent transform (Fisher, 1921) at the subject level to the beta-series correlation values (r values) of each pair of ROIs and each experimental condition. Since the correlation coefficient is inherently restricted to range from −1 to +1, this transformation ensured the null hypothesis sampling distribution approached that of the normal distribution. First, we created correlation matrices with the mean correlation values of all subjects for each experimental condition, where significant correlations, r ≥ 0.57 were colored red. In this case, Pearson’s r values were corrected using Bonferroni correction (Pair of ROIs x Conditions x Hemisphere), which is more conservative than FDR, since we were testing coactivation between adjacent regions. Afterwards, we examined the association between functional connectivity strength and physical distance (i.e., the Euclidean distance between the center of mass of two ROIs) of the given ROI pair per subject. The Euclidean distance between the center of mass of two ROIs was computed by averaging the left and right hemisphere Euclidean distance values. Finally, to test for significant differences in the coupling strength of the pairwise connectivity, Fisher’s Z normally distributed values for each pair of ROIs for each participant and condition were submitted to 6 separated 5 (ROI Pair) x 2 (Pathway) x 2 (Stimuli) repeated measure ANOVAs. Results were corrected using the Bonferroni correction for multiple comparisons.

#### Laterality analyses

Lateralization indexes of the signal t-values for the four conditions of interest, i.e., magnocellular-biased words, magnocellular-biased images, parvocellular-biased words and parvocellular-biased images were calculated. For this purpose, the standard lateralization index (LI) was computed:

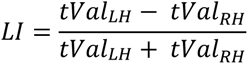

This index produces values between -1 and 1 and relates the left-right signal t-value difference to the mean of the left-right signal t-values. Positive values correspond to stronger left than right t-values; conversely, negative values correspond to stronger right than left t- values.

Since LI values must be between 0 and 1, the t-values were first normalized to values from 0 and 1 using a linear scaling procedure. Once signal t-values were normalized, the laterality of words and images was computed for the magnocellular and parvocellular pathways within the visual regions. Then, paired t-tests between words and images LIs were performed for each of the pathways and ROIs. Results were corrected using the false discovery rate (FDR) correction for multiple comparisons. The laterality of two pathways was not compared since the ROI analysis, probabilistic maps, and functional connectivity analysis did not reveal relevant significant effects of hemisphere as a function of pathway.

Finally, to test the reproducibility of our results, 14 participants were invited to a retest session and underwent the exact same MRI protocol and functional tasks 7-10 days after they had completed the initial session. ROI, probabilistic, and functional connectivity results for this retest session are shown in Figure 3, 4 and 5 supplement 2.

## 3. Results

### 3.1 In-scanner behavioral results

A 2 (Pathway) x 2 (Stimuli) ANOVA revealed a statistically significant main effect of Pathway (F_1,31_= 12.84; ηp^2^ =0.29, p < 0.001, BF10 = 9.91), (see Fig. 2*A*), showing that accuracy was higher for parvocellular-biased than magnocellular-biased stimuli.

**Figure 2.**
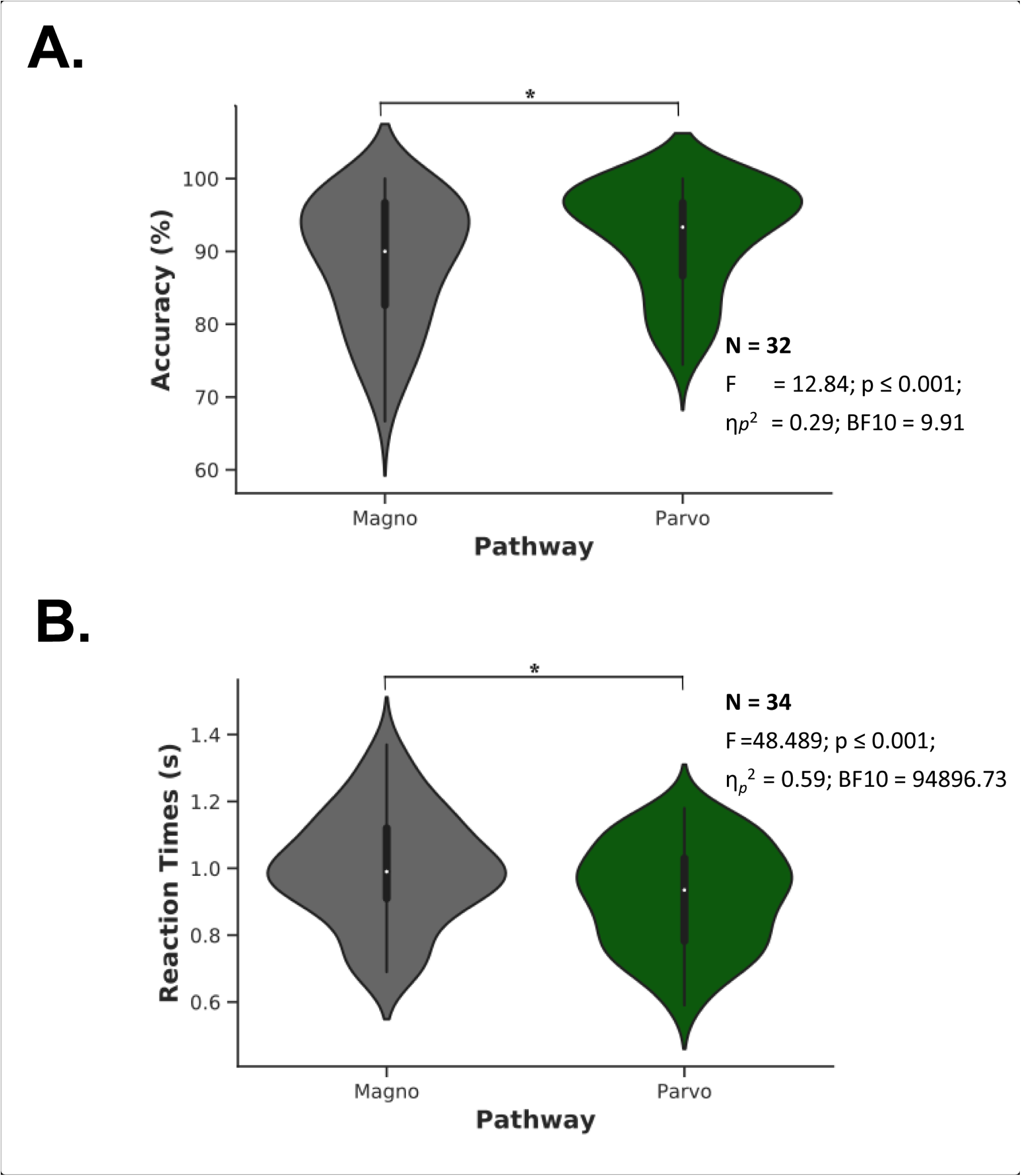
In-scanner behavioral results. A) Percent (%) mean accuracy of manmade and non- manmade judgments as a function of visual magnocellular and parvocellular pathways. B) Correct-response reaction times as a function of magnocellular and parvocellular pathways. The white dots on the violin plots represents the median, while the black bar in the center represents the interquartile range. Thick black lines indicate statistically significant effects: *ps < .05.

The ANOVA for reaction times to correct responses also revealed significant main effects of Pathway (F_1,33_=48.489; p ≤ 0.001, η*_p_*^2^ =0.595, BF10 = 94896.73) and Stimuli (F_1,33_=33.30; p ≤ 0.001, η*_p_*^2^ =0.50, BF10 = 106557.38). Simple-effect *post-hoc* analyses revealed the main effect of pathway was due to longer response latencies for magnocellular-biased than parvocellular-biased stimuli (see Fig. 2*B*). The main effect of Stimuli was driven by longer response times for words (M = 1.00 s; SD = 0.16 s) than images (M = 0.92 s; SD = 0.15 s). The Pathway x Stimuli interaction was not significant (F_1,33_=2.18; p = 0.149, η*_p_*^2^ =0.06, BF10 = 1.62 e^12^).

### 3.2 fMRI results

We used four analytic approaches to examine visual recognition of words and images in the visual cortex and the degree of involvement of the two main magnocellular and parvocellular visual pathways. All analyses were conducted in individual-subject space. First, given that the activation signal tends to be stronger in the more posterior relative to the more anterior visual areas, we used specific contrasts in ROI-related analyses, including both percent signal change and signal t-value dependent measures, to examine the recruitment of visual cortex regions in line with our experimental design. Second, to further investigate differential contributions of visual regions to the two main visual pathways for reading and image recognition, we created probabilistic maps using the same contrasts utilized in the ROI analyses. In our third analytical approach, we used functional connectivity methods to examine differential coactivation among visual regions during task processing. We computed the relationship between the pairwise-connectivity correlation values and the Euclidean distance between the physical distances of the center of mass of each of the ROI pairs, along with analyses comparing the strength of coactivations between pairs of regions. Finally, we investigated lateralization effects associated with word and image processing specifically for each visual pathway given the lack of differences between hemispheres for the pathway factor in our previous analyses.

All these analytical steps, except the laterality analysis, were conducted for left-lateralized visual regions because our focus was on language-related processes. Nevertheless, we also present the corresponding right-hemisphere analyses in Figure 3, 4 and 5 supplement 1.

#### 3.2.1 Individual ROI analyses

ROI analyses were conducted at the individual-subject level to characterize the activation profile of regions within the left visual cortex. To avoid potential biases in the observed effects due to stronger signal strength in more posterior relative to more anterior regions, parameter estimates (i.e., scaled % signal change and signal t-values) for the specific contrasts (Magnocellular > Parvocellular Words and Magnocellular > Parvocellular Images) were extracted and submitted to a 6 (ROI: V1, V2, V3, hV4, LO1, TO1) X 2 (Stimuli: words, images) repeated measures ANOVA. This analysis revealed main effects of ROI (F_5,150_ = 50.01; p ≤ 0.001, η*_p_*^2^ = 0.62, BF10 = 2.38 e^31^) and Stimuli (F_1,30_ = 1.21; p = 0.006, η*_p_*^2^ = 0.25, BF10 = 547.25). Simple-effect analyses for the ROI main effect revealed an anterior to posterior functional gradient with stronger engagement of the more anterior relative to the most posterior visual cortical regions (see Fig. 3*A*, upper panel). All comparisons were statistically significant (ps ≤ 0.001, d’ ≥ 852, BF10 ≥ 120170.861; V2-hV4 and LO1-TO1 ps ≤ 0.042, d’ ≥ 0.331, BF10 ≥ 2.788), with the exception of hV4-V3, the only comparison that did not show a statistical difference (p = 0.385, d’ = 0.089, BF10 = 0.177). In regard to the main effect of Stimuli, these regions in visual cortex were in general more strongly engaged by processing words (M = -0.11; SD = 0.32) than images (M = -0.20; SD = 0.33).

**Figure 3.**
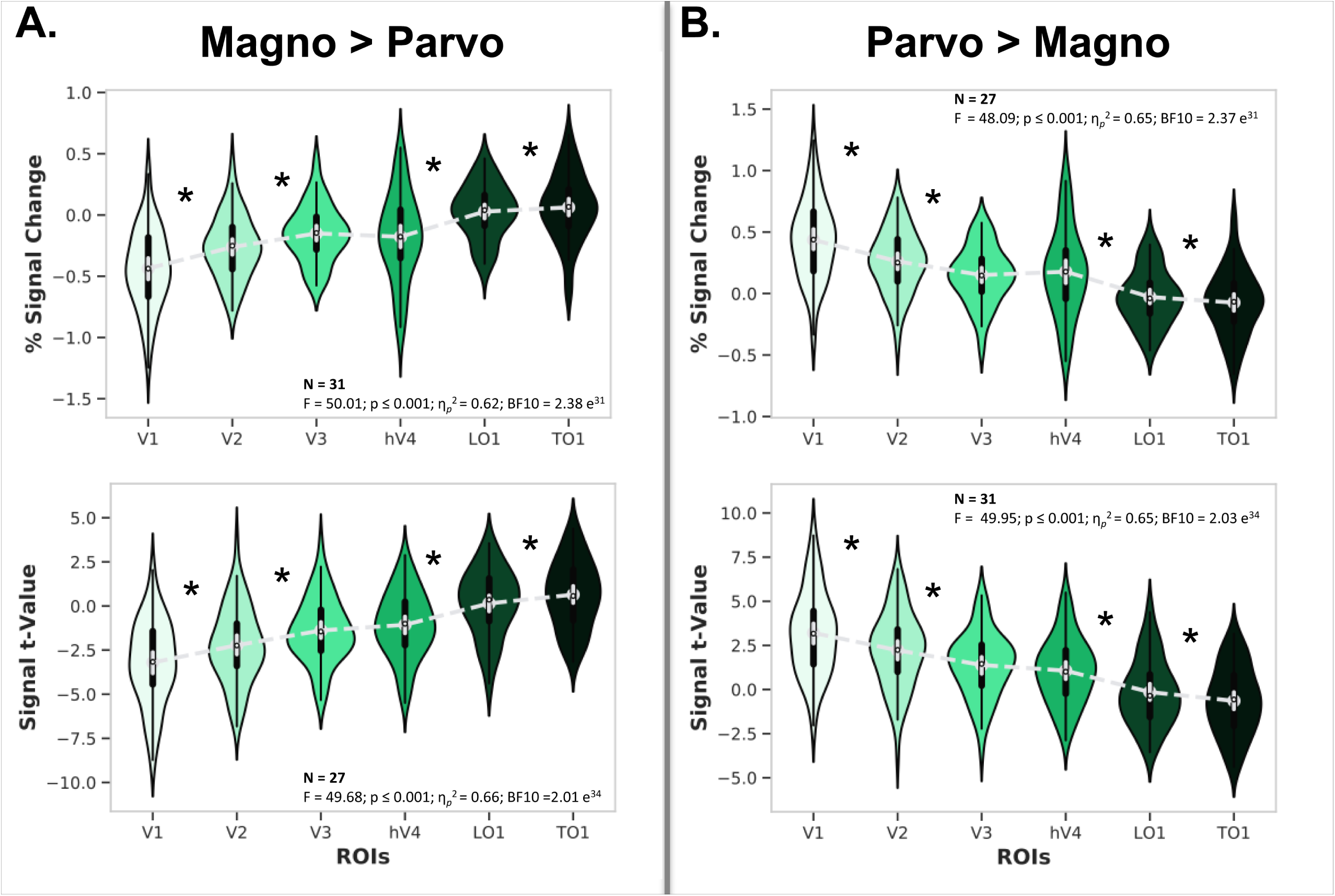
ROI analyses. A) Percent signal change (top panel) and regional t-values (bottom panel) of left hemisphere visual cortex ROIs for the Magnocellular > Parvocellular contrast. B) Percent signal change (top panel) and regional t-values (bottom panel) of left hemisphere visual cortex ROIs for the Parvocellular > Magnocellular contrast. The black bar in the center of the violins represents the interquartile range, while the gray dash lines reflect the median. * asterisks denote statistically significant effects (FDR corrected). Magno = magnocellular; Parvo = parvocellular.

Second, the same analysis was carried out with t-values instead of % signal change values. The same general pattern of results emerged (ROI main effect, F_5,130_ = 49.68; p ≤ 0.001, η*_p_*^2^ = 0.66, BF10 =2.01 e^34^), except that main effect of Stimuli did not reach significance (F_1,26_ = 3.04; p = 0.093, η*_p_*^2^ = 0.11, BF10 = 2.47). Simple-effect analysis for the ROI main effect also confirmed an anterior-posterior functional gradient for the scaled % signal change values. All comparisons were statistically significant (ps ≤ 0.001, d’ ≥ 0.821, BF10 ≥ 184534.827; V2-hV4, p = 0.042, d’ = 0.704, BF10 =12697.017; LO1-TO1, p = 0.013, d’ = 0.393, BF10 = 9.339), except the hV4-V3 comparison, which revealed no statistical difference (p = 0.167, d’ = 0.260, BF10 =0.887). Figure 3 shows the ROI analysis for Magnocellular > Parvocellular contrasts across stimuli conditions, given that the interaction ROI x Stimuli was not significant (Fig 3*A*: the upper panel corresponds to % signal change values; the bottom panel corresponds to signal t-values).

Third, the same 6 X 2 repeated measures ANOVAs using % signal change and t-values as dependent measures was conducted for the specific contrasts Parvocellular > Magnocellular Words and Parvocellular > Magnocellular Images for each left visual cortex ROIs. Again, the same pattern of results emerged in the ANOVAs and post-hoc analyses. % signal change (Fig. 3*B*, upper panel) showed main effects of ROI (F_5,130_ = 48.09; p ≤ 0.001, η*_p_*^2^ = 0.65, BF10 = 2.37 e^31^) and Stimuli (F_1,26_ = 9.00; p = 0.006, η*_p_*^2^ = 0.26), while t-values (Fig. 3*B*, bottom panel) showed a main effect of ROI (F_5,135_ = 49.75; p ≤ 0.001, η*_p_*^2^ = 0.65, BF10 = 2.03 e^34^).

Fourth, identical ROI analyses were carried out for the right visual cortex (see Figure 3- supplement 1). Findings were similar to those observed for left visual cortex in terms of the anterior-posterior functional gradient (i.e., ROI main effect). However, in contrast to the left hemisphere where we observed a main effect of Stimuli for % signal change values (but not signal t-values), the right hemisphere analysis did not show a main effect of Stimuli in either the % signal change or signal t-values dependent measures. This suggests that the left hemisphere might be more sensitive in differentiating the two types of stimuli (words and images), but only when parameter estimates are used as the dependent measure.

Finally, identical ROI analyses were conducted for the left and right visual cortex (see Figure 3- supplement 2) in the retest session with 14 participants. Findings were similar to those observed in the main experiment in terms of the anterior-posterior functional gradient (i.e., ROI main effect).

#### 3.2.2 Probabilistic maps

To further examining the robustness of the functional gradient observed for the Pathway factor in terms of % signal change and signal t-values, we conducted an additional fMRI analysis using left visual cortex surface-based probabilistic maps. This allowed us to examine the probability that a given voxel was significantly engaged in a functional contrast of interest. Since we did not observe any significant interactions with the factor Stimuli in the ROI Analysis, we performed probabilistic maps for Magnocellular > Parvocellular and Parvocellular > Magnocellular contrasts across stimuli conditions. These maps revealed that in the Magnocellular > Parvocellular contrast, higher probabilities were located in more anterior regions within the visual cortex (i.e., LO1, TO1; see Fig. 4*A*). In contrast, the probabilistic map for Parvocellular > Magnocellular showed higher probabilities in more posterior regions, including primary visual cortex (V1, V2, V3 and hV4; Fig. 4*B*). Color-coding denotes the likelihood of each voxel being assigned to one or the other pathway in the probabilistic map. Maps are not fully complementary since probability is assigned based on each specific contrast.

**Figure 4.**
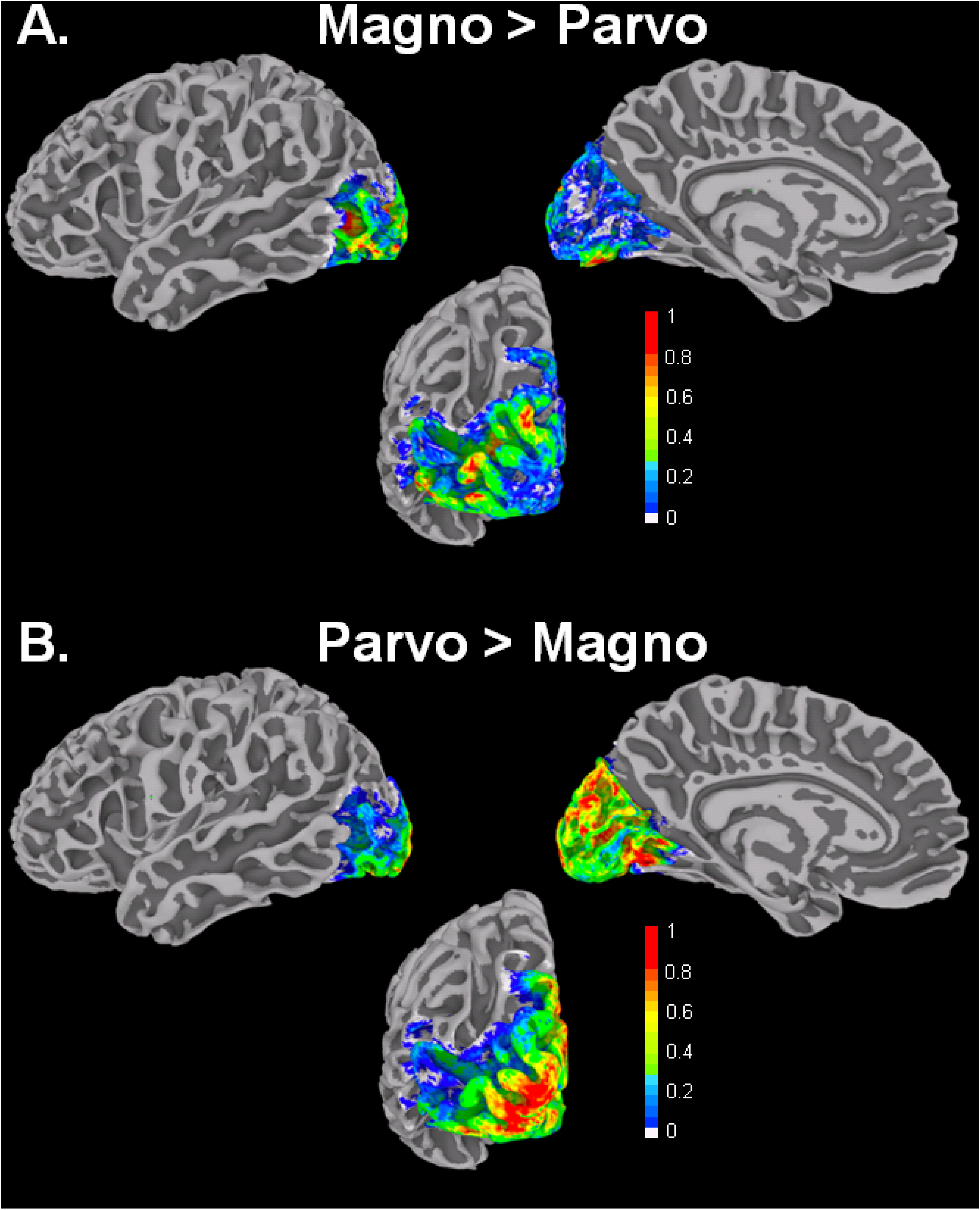
Probabilistic maps showing the percentage of involvement of left visual cortex voxels on surface renderings for the contrasts A) Magno > Parvo and B) Parvo > Magno. The two probabilistic maps are not fully complementary: although they represent opposite contrasts, they were generated by separate analyses. To create these probabilistic maps activations inside the visual cortex ROIs were binarized for the two contrasts by using a threshold common to all the subjects; for every subject, all voxels that had a positive t-values were scored 1 and used for the magnocellular-biased maps with the rest zeroed, whereas all voxels that had a negative t-values were scored 1 and used for the parvocellular-biased maps with the rest zeroed. Magno = magnocellular; Parvo = parvocellular.

Probabilistic maps for the right visual cortex showed a similar pattern to those for the left hemisphere, with stronger probabilities for the involvement of anterior regions in the Magnocellular > Parvocellular contrast and for posterior regions in the Parvocellular > Magnocellular contrast (see Figure 4- supplement 1).

Finally, probabilistic maps for the left and right visual cortex from the retest session also revealed a similar pattern to those for the left and right hemisphere in the main experiment, with stronger probabilities for the involvement of anterior regions in the Magnocellular > Parvocellular contrast and for posterior regions in the Parvocellular > Magnocellular contrast (see Figure 4- supplement 2).

#### 3.2.3 Pairwise functional connectivity analysis

To examine patterns of functional connectivity between left visual cortex regions, we first characterized overall coupling strength across study conditions. Figure 5*A* shows the correlation matrix for Pearson’s r values between pairs of ROIs (red colored circles indicate statistically significant FC values, r ≥ 0.57, Bonferroni corrected) and a representation of Euclidean distance against Pearson’s r values between each pair of nodes (shadowed areas indicate significant FC). Figure 5*B* shows a surface rendering with statistically significant functional connectivity between visual cortex regions displayed as edges. Overall, stronger coactivations were found among posterior visual regions (i.e., V1, V2, V3) and anterior visual regions (i.e., hV4, TO1, TO1). In contrast, more distant anterior-posterior regions did not show statistically significant FC.

**Figure 5.**
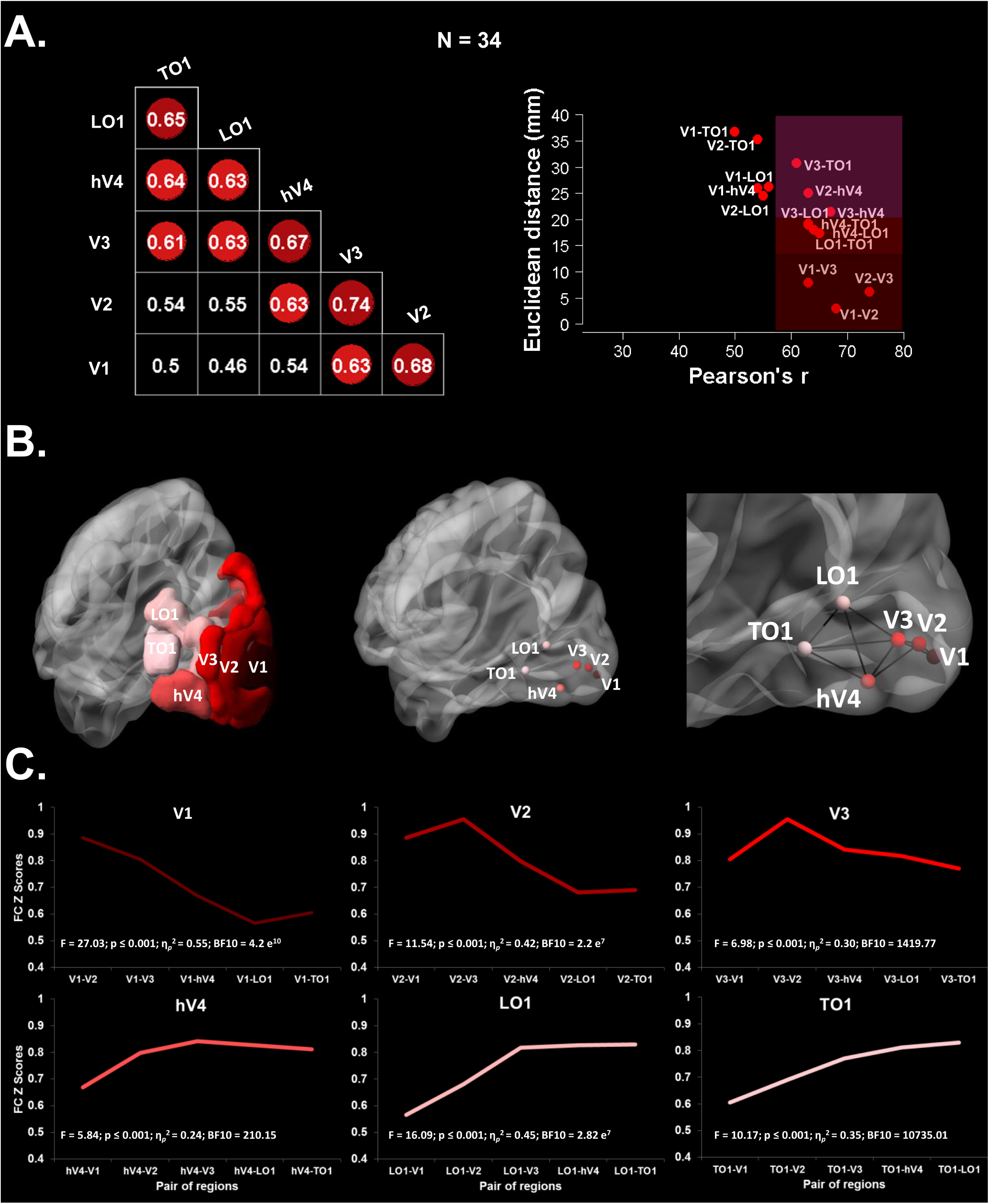
Task-related pairwise functional connectivity between left visual cortex ROIs across experimental conditions (i.e., magnocellular-biased words, parvocellular-biased words, magnocellular-biased images, parvocellular-biased images). A) Left panel shows the correlation matrix with significant Pearson’s r correlations (r ≥. 57, Bonferroni corrected) colored in red tones. Right panel shows Euclidean distances from posterior (V1, 0-value) to anterior (TO1) left visual cortex regions plotted against Pearson’s r values for each ROI pair; shaded areas indicate pairs with significant coupling (r ≥. 57, Bonferroni corrected). B) Left panel shows the sagittal rendering of the left visual cortex ROIs. Central panel shows the sagittal rendering of left visual cortex nodes centered at the ROIs center of mass. Right panel shows statistically significant pairwise correlations (i.e., edges) between left visual cortex nodes in a posterior sagittal rendering section (r ≥. 57, Bonferroni corrected). C) Task-related pairwise functional connectivity (Z scores) between left visual cortex ROI pairs across experimental conditions. The upper left panel shows Z scores between V1 and the rest of the ROIs, upper middle panel shows Z scores between V2 and the rest of the ROIs, and upper right panel shows Z scores between V3 and the rest of the ROIs. The lower left panel shows Z scores between hV4 and the rest of the ROIs, lower middle panel shows Z scores between LO1 and the rest of the ROIs, and lower right panel shows Z scores between TO1 and the rest of the ROIs.

Second, to more closely examine FC in the left visual cortex, in line with our experimental questions, Fisher’s Z transformed FC values were submitted to a series of 5 (ROI Pairs) x 2 (Pathway) x 2 (Stimuli) separate repeated measures ANOVAs, one for each of the examined visual cortex regions. All of the analyses revealed a main effect only for ROI Pair (F_4,132_ ≥ 5.84; ps ≤ 0.001, η*_p_*^2^ ≥ 0.24, BF10 ≥ 210.15) (Fig. 5*C*). From more posterior to more anterior, simple-effect analyses revealed that *V1* was functionally more strongly tied to V2 than to V3, hV4, LO1 and TO1 (ps ≤ 0.04, d’ ≥ 0.411, BF10 ≥ 40360.043); but no differences in FC emerged between V1-LO1 or V1-TO1 or between V1-hV4 or V1-TO1 (ps ≥ 0.05, d’ ≤ 0.263, BF10 ≤ 6.392; see Fig. 5*C*). Second, there was stronger FC between *V2* and V3 than between V2 and all the other visual regions (i.e., V1, V3, hV4, LO1 and TO1; ps ≤ 0.04, d’ ≥ 0.270, BF10 ≥ 6.615); again, with no significant differences emerging in FC between V2-LO1 and V2-TO1 (ps ≥ 0.05, d’ = 0.024, BF10 = 0.104). Third, a similar pattern of results emerged for *V3*: stronger FC for the V3-V2 contrast than for contrasts between V3 and the rest of the visual regions (i.e., V1, hV4, LO1 and TO1; ps ≤ 0.02, d’ ≥ 0.280, BF10 ≥ 9.844; no FC differences between V3-LO1 and V3-TO1, V3-V1 and V3-hV4, V3-V1 and V3-LO1, or V3-V1 and V3-TO1 [ps ≥ 0.05, d’ ≤ 0.162, BF10 ≤ 0.474]).

On the other hand, more anterior visual cortex regions hV4, LO1 and TO1 presented a similar FC pattern. hV4 showed stronger connectivity with V3 relative to either V2 or V1 (ps ≤ 0.01, d’ ≥ 0.314, BF10 ≥ 31.706), but there were no differences in coactivation between hV4-V1 and hV4-LO1, hV4-V3 and hV4-LO1, hV4-V2 and hV4-LO1, hV4-V2 and hV4- TO1, hV4-V3 and hV4-TO1, or hV4-LO1 and hV4-TO1 (ps ≥ 0.05, d’ ≤ 0.103, BF10 ≤ 0.189; see Fig. 5*C*). FC of the *LO1* was similar to visual regions V3, hV4, and TO1; and stronger to all of these regions than to visual cortex regions V1 and V2 (ps ≤ 0.04, d’ ≥ 0.376, BF10 ≥ 265.743); but there were no differences in coactivation, between LO1-V3 and LO1- hV4, LO1-V3 and LO1-TO1, or between LO1-hV4 and LO1-TO1 (ps ≥ 0.05, d’ ≤ 0.060, BF10 ≤ 0.124). Finally, FC of the *TO1* was similar to that for visual regions TO1 and hV4; exhibiting stronger connections to these two regions compared to visual cortex regions V1, V2, and V3 (ps ≤ 0.01, d’ ≥ 0.293, BF10 ≥ 265.743). There were no differences in the coactivation between TO1-V3 and TO1-hV4, TO1-V3, or TO1-LO1 or between TO1-hV4 and TO1-LO1 (ps ≥ 0.05, d’ ≤ 0.216, BF10 ≥ 0.124).

In sum, V1, V2, and V3 were tightly coupled to each other as a function of their proximity, showing statistically significant lower FC with the more anterior visual cortex regions (LO1, TO1) (Fig. 5*C*). Anterior regions hV4, LO1, and TO1 were also strongly coupled to each other but showed statistically lower FC with posterior regions V1 and V2. FC results for the right hemisphere are shown in Figure 5- supplement 1. FC results from the retests session for the left and right hemisphere are shown in Figure 5- supplement 2. Results from all these analyses revealed a similar pattern as those described for the left hemisphere.

#### 3.2.4 Laterality analysis

To study hemispheric specialization, paired t-tests were conducted on laterality indexes of words and images for each ROI within each pathway. Laterality analyses comparing the two pathways were not performed since ROI analysis, probability maps, and FC analysis did not reveal relevant significant hemispheric effects as a function of pathway. Comparisons between magnocellular-biased words and images revealed that the LO1 region was more left lateralized for words (LI = 0.09) than images (LI = -0.16) (p = 0.003, d’ = 0.68, BF10 = 63.83). For the parvocellular pathway, results showed that the V1 region was also more left lateralized for images (LI = 0.02) than words (LI = -0.10) (p = 0.022, d’ = 0.52, BF10 = 7.12). All the other comparisons did not reveal any statistically significant effects (ps ≥ 0.053, d’ ≤ 0.44, BF10 ≤ 2.92).

## 4. Discussion

In the current study, visual cortex magnocellular and parvocellular contributions to reading and object recognition were examined. Although there are well-accepted theories regarding the roles of these main visual pathways in object recognition, their functional involvement of the in word recognition has received considerably less attention. The present study sought to address this gap in the research by examining the functional correlates of word and image recognition along these pathways within the visual cortex. Our analytical approach included behavioral analyses and both regional and connectivity functional MRI measures in individual-subject space. Results showed (1) an advantage for parvocellular-biased over magnocellular-biased stimuli in terms of accuracy and reaction times; (2) differential recruitment of posterior versus anterior visual cortex regions as a function of the parvocellular versus magnocellular pathway manipulation in our experimental design; (3) functional coupling as a function of the distance between regions, with more clustering within than between posterior and anterior regions (4) more left lateralization of words in more peripheral regions of the visual cortex, but more left lateralization of images in more posterior regions of the visual cortex. Results were extensively replicated in a retest session carried out with a subset of participants (n =14) 7-10 days after the initial session (see Figure 3, 4 and 5 supplement 2). The relevance of these findings for reading is discussed next.

Magnocellular-biased stimuli are less bright and more difficult to recognize than parvocellular-biased stimuli. This led to lower accuracy and longer reaction times for these stimuli than either the parvocellular-biased or neutral non-biased stimuli in the behavioral results – even though we had conducted a pilot study to ensure high accuracy (i.e., 80%) and a sufficient number of correct answers for the magnocellular-biased stimuli.

Regional activation profiles and probabilistic maps revealed different functional patterns of activation within the visual cortex, along a gradient across the regions of the visual cortex, with parvocellular-biased stimuli recruiting posterior visual cortex regions more strongly than magnocellular-biased stimuli, and magnocellular-biased stimuli engaging anterior visual cortex regions more strongly than parvocellular-biased stimuli. Previous evidence from monkeys has revealed that posterior visual cortex regions (V1, V2, and V3) respond to both magnocellular-biased and parvocellular-biased stimuli. Other studies in humans, such as Musel et al. (2013), have observed that low spatial frequencies (associated with the magnocellular pathway) activate V1, whereas high spatial frequencies (associated with the parvocellular pathway) activate V2, V3. Here we found that areas V1, V2, V3 all showed a preference for parvocellular- over magnocellular-biased stimuli. However, unlike Musel et al. (2013), our stimuli were designed to take all the properties associated with the magnocellular and parvocellular bias into account, not only spatial frequency. This may have led to the different patterns of activation observed in the two studies.

Following the gradient of the visual cortex and in line with previous findings, the hV4 region was also more strongly recruited by parvocellular- than magnocellular-biased stimuli. This result is highly consistent with previous neuroimaging studies with both healthy and clinical patients, which demonstrated that the hV4 region is in charge of color processing (Pearlman et al., 1979, Lueck et al. 1989, Zeki et al., 1990, McKeefry & Zeki 1997, Bartels and Zeki, 2000). Similarly, previous visual recognition studies have suggested that hV4 forms part of the ventral visual stream, which is mainly associated with object, feature, and form processing (Bartels and Zeki, 2000; Brewer et al., 2005; Arcaro et al., 2009).

In contrast, anterior visual cortex areas LO1 and TO1 showed stronger activation for magnocellular- than parvocellular-biased stimuli (Sayres et al., 2008; Skottun et al., 2015, Stigliani et al., 2017). MT+/V5 region, part of TO1, is a region with high contrast sensitivity that processes motion, and is therofre activated more by moving than stationary stimuli. Tootell et al. (1995) showed that MT+ contains direction-selective neural populations, while as Huk et al. (2002) demonstrated that this region contains pattern-motion cells. Although the LO complex has always been seen as an object processing region, Larsson et al., (2006) found that LO1 responds to stimuli designed to test orientation selectivity, a property that is associated with magnocellular pathway processes.

Our results represent solid and converging evidence for a posterior to anterior functional gradient within the visual cortex for processing parvocellular- and magnocellular-biased stimuli. Further, we have shown that this effect holds regardless of the nature of the stimuli (words or images), and is largely similar across the left and right hemispheres. These findings hold considerable promise for further research aimed at unraveling the contributions of the visual cortex to reading processes. Future studies should examine to what extent the contributions of the magnocellular and parvocellular systems to reading and visual recognition change over development, especially after reading acquisition, and examine potential differences in these functional gradients in typical and atypical readers (Müller-Axt et al., 2017).

Pairwise functional connectivity analyses revealed differential functional coupling as a function of physical distance between pairs of ROIs, regardless of the visual pathway or stimuli type. We observed significant functional connections between regions within the posterior or early visual cortex (i.e., V1, V2, V3) and between regions in the more anterior and peripheral regions (LO1 and TO1). Region hV4 was an exception, exhibiting an intermediate pattern of activation, with tight coupling to both more posterior and more anterior regions.

Furthermore, we demonstrated that pairwise functional connectivity between regions in the visual cortex is closely related to their physical distance, with decreasing functional connectivity coefficients for greater physical distances. This was established by our analysis of correlations between the functional connectivity and physical distance between pairs of regions as well as by ANOVA analyses of Z-scores for functional connectivity (ROI Pair x Pathway x Stimuli). Studies using resting-state functional connectivity have previously reported that physical distance and connectivity measures are strongly related (Genç et al., 2015, Dawson et al., 2016).

Turning to the question of hemispheric lateralization, in line with our hypothesis, we found that word stimuli were left-lateralized in more anterior regions, specifically LO1, for magnocellular-biased stimuli, while image stimuli were left-lateralized in V1 for parvocellular-biased stimuli. Recent neuroimaging studies have suggested that there is a left lateralized region involved in word recognition, which they have called the Occipital Word Form Area (OWFA) (Yu et al., 2015, Strother et al., 2015, Strother et al., 2017). For instance, Strother et al. (2017) reported that the OWFA is located intermediate to hV4, and within the lateral occipital gyrus (LO). Furthermore, in line with our results, Brederoo et al.’s (2017) study found left lateralization for global linguistic and non-linguistic stimuli (global processing can be associated with the magnocellular pathway). Nevertheless, contrary to our results, the same study found a right hemisphere preference for non-linguistic stimuli, whereas we found left-hemisphere preference for images in the primary visual cortex. This may be due to differences in the stimuli and tasks used in the two studies. While we used line drawings as images that were chromatically defined and isoluminant, Brederoo et al. (2017) used compound stimuli with targets made up of black diamonds or plus signs and distractors made up of rectangles or crosses, all displayed against white backgrounds, following classical Navon (1977) task procedures.

In sum, in our study we found a functional gradient in the activation of the visual cortex, where posterior visual regions were more strongly recruited for parvocellular-biased stimuli, whereas more anterior regions were more engaged by magnocellular-biased stimuli. Functional connectivity between visual regions depended on physical distance and appeared to be segregated, with the strongest FC observed as clusters within these more posterior and anterior regions. Finally, we saw that left-lateralization for words occurred in more anterior regions for magnocellular-biased stimuli but in more posterior regions for images. Altogether, these findings provide converging evidence for functional division of labor in the visual cortex as a function of the parvocellular and magnocellular properties of the stimuli.

## Acknowledgments

We thank Kestas Kveraga for providing us with initial versions of the scripts used to produce magnocellular and parvocellular stimuli; Asier Zarraga and David Soto for technical support; Jon Imanol Etxabe for helping with data collection; and Magda Altman for help editing the manuscript. This work was supported by MINECO predoctoral grants from the Spanish government (BES-2016-077568) to M.I.-G.; grants from the Ministerio de Ciencia e Innovación (RYC-2014-15440; PSI2015-65696; PGC2018-093408-B-I00), Neuroscience projects from the Fundación Tatiana Pérez de Guzmán el Bueno, Basque Government (PIBA- 2021-1-16-0003), and “la Caixa” Banking Foundation under the project code LCF/PR/HR19/52160002 to P.M.P.-A. BCBL acknowledges support from the Basque Government through the BERC 2022-2025 program and from the Spanish State Research Agency through BCBL’s Severo Ochoa excellence accreditation CEX2020-001010-S.

## 5. Author contributions

M.I.-G. and P.M.P-A conceived and designed the study, performed the experiments, analyzed the data and wrote the manuscript.

---

**Figure 3- supplement 1.**
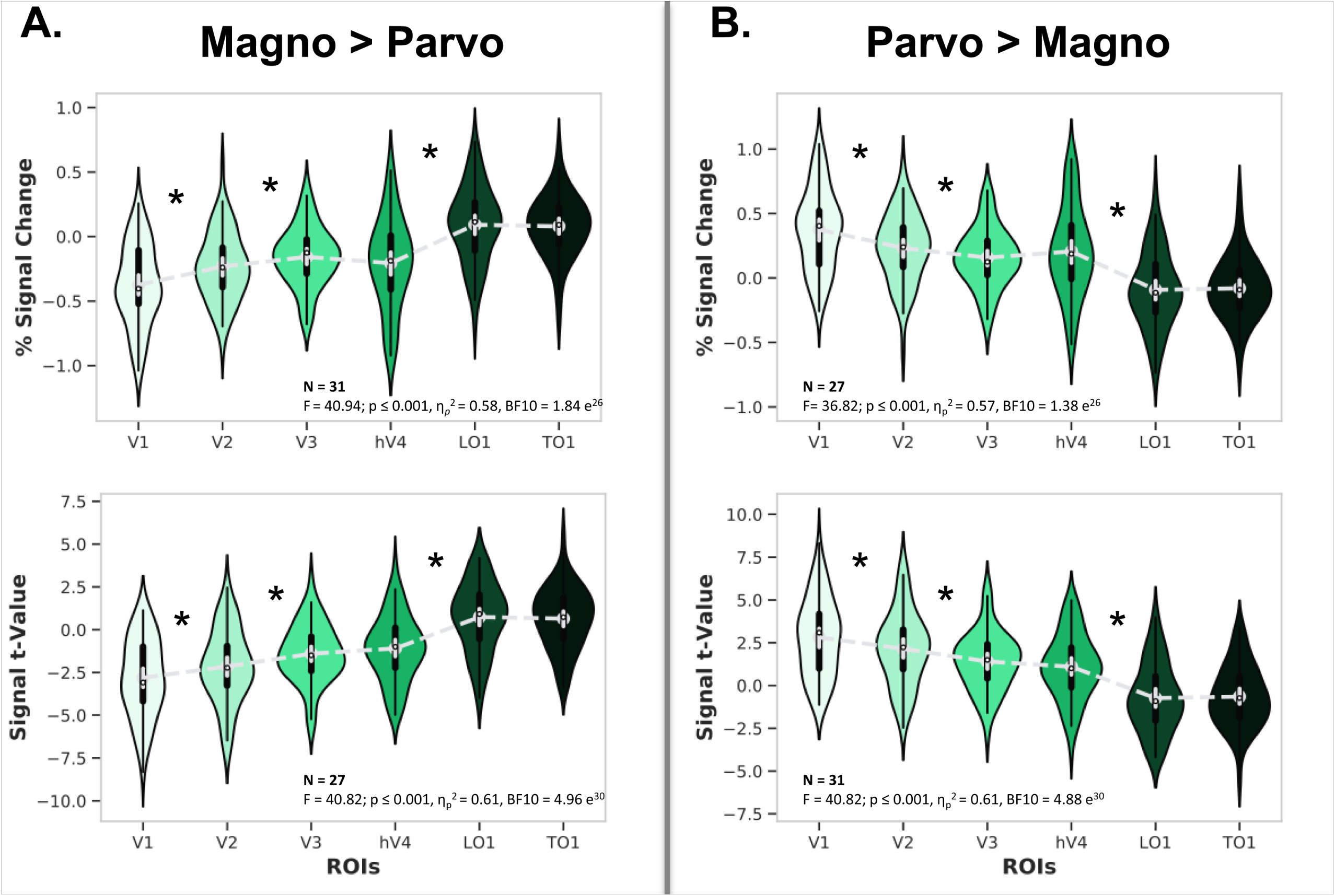
Right hemisphere ROI analyses. A) Percent signal change (top panel) and regional t-values (bottom panel) of right hemisphere visual cortex ROIs for the Magnocellular > Parvocellular contrast. B) Percent signal change (top panel) and regional t-values (bottom panel) of right hemisphere visual cortex ROIs for the Parvocellular > Magnocellular contrast. The black bar in the center of the violins represents the interquartile range, while the gray dash lines reflect the median. * asterisks denote statistically significant effects between contiguous regions (FDR corrected). Magno = magnocellular; Parvo = parvocellular.

**Figure 3- supplement 2.**
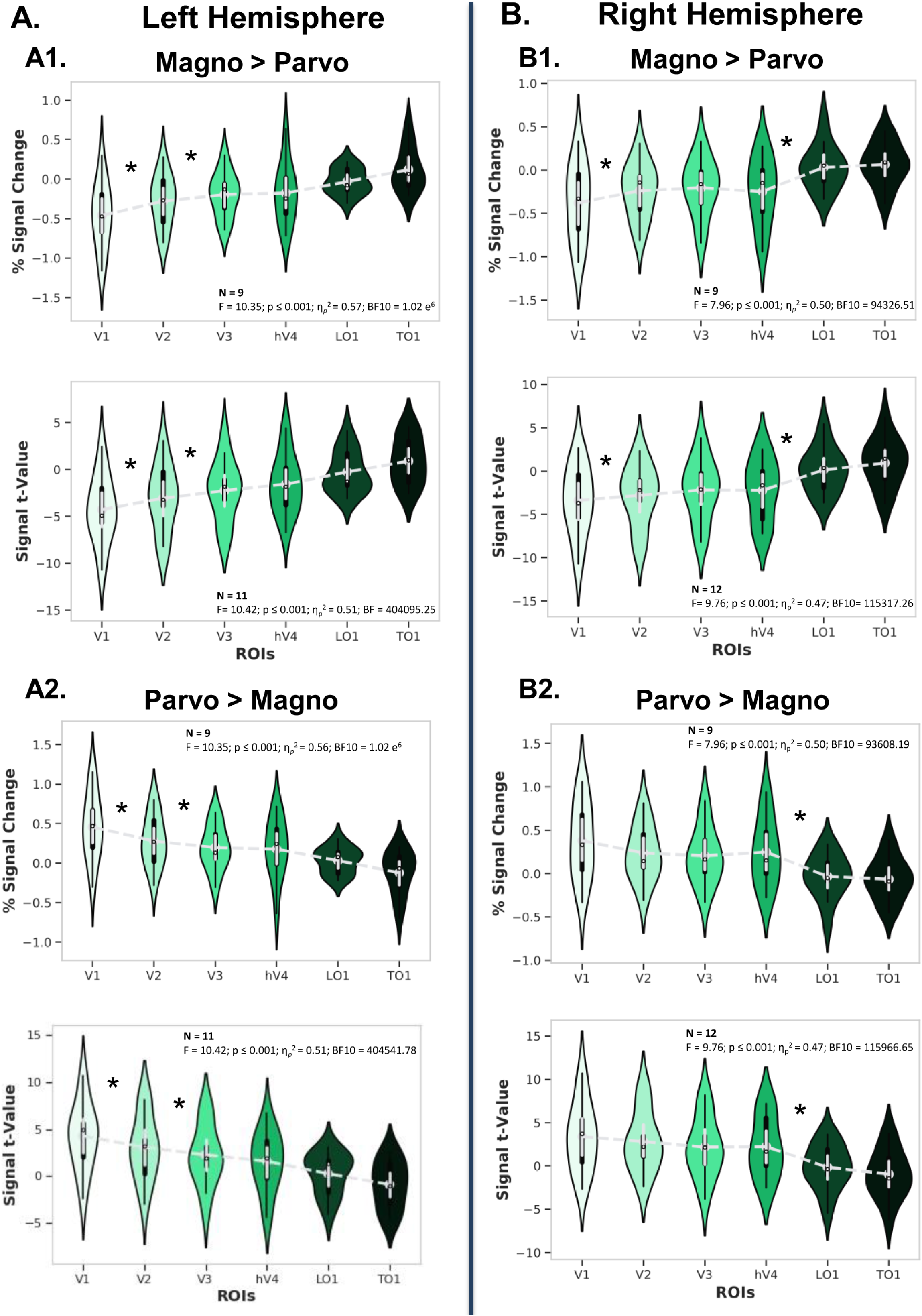
Retest session left and right hemisphere ROI analyses. A) Left hemisphere ROI analysis, with A.1) showing percent signal change (top panel) and regional t-values (bottom panel) of visual cortex ROIs for the Magnocellular > Parvocellular contrast and with A.2) showing percent signal change (top panel) and regional t-values (bottom panel) of visual cortex ROIs for the Parvocellular > Magnocellular contrast. B) Right hemisphere ROI analysis, with B.1) showing percent signal change (top panel) and regional t-values (bottom panel) of visual cortex ROIs for the Magnocellular > Parvocellular contrast and with B.2) showing percent signal change (top panel) and regional t-values (bottom panel) of visual cortex ROIs for the Parvocellular > Magnocellular contrast. The black bars in the center of the violins represents the interquartile range, while the gray dash lines reflect the median. * asterisks denote statistically significant effects between contiguous regions (FDR corrected). Magno = magnocellular; Parvo = parvocellular.

**Figure 4- supplement 1.**
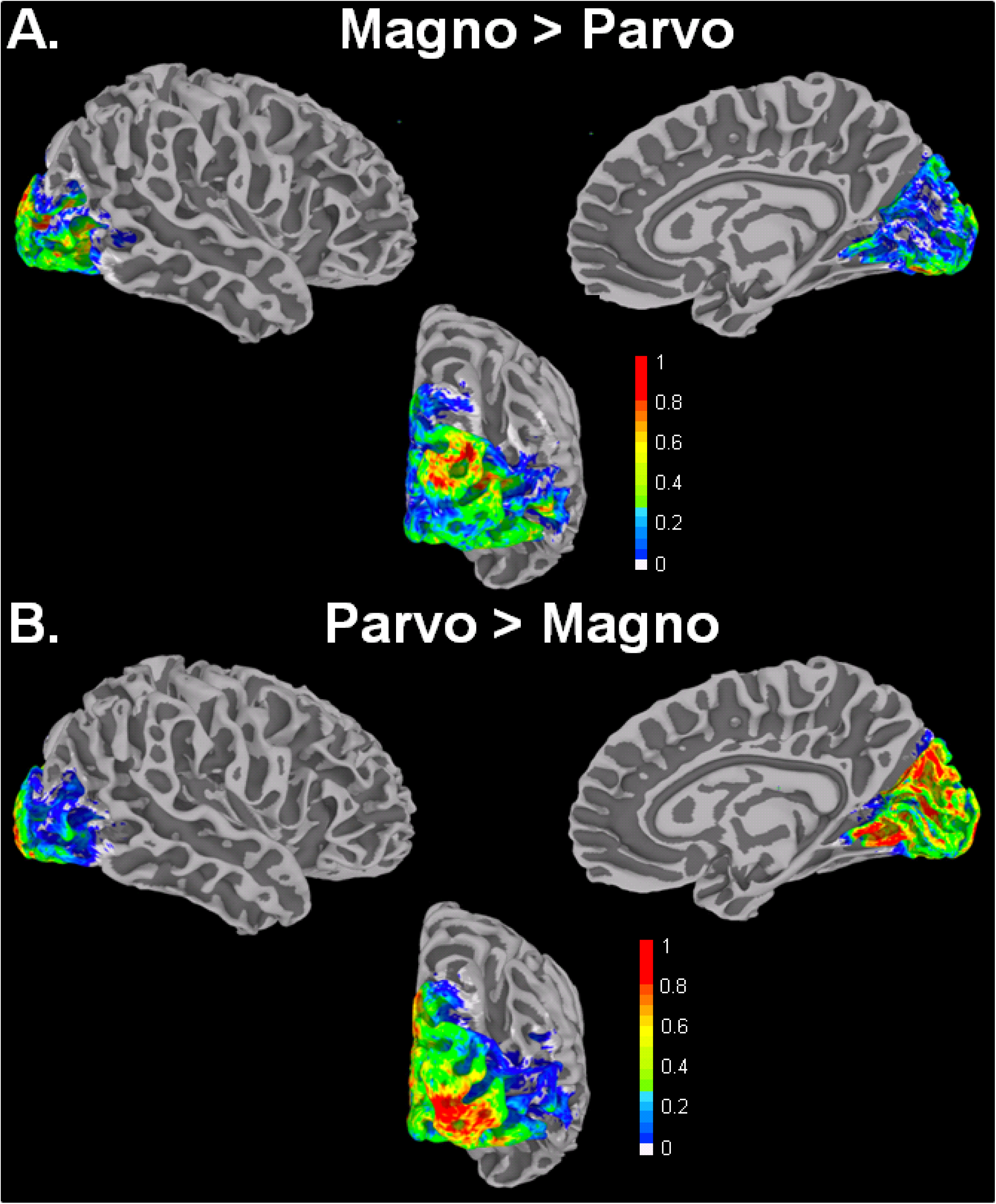
Right hemisphere probabilistic maps showing the percentage of involvement of visual cortex voxels on surface renderings for the contrasts A) Magno > Parvo and B) Parvo > Magno. The two probabilistic maps are not fully complementary in terms of voxel correspondence; while they represent opposite contrasts, they were generated by separate analyses. Magno = magnocellular; Parvo = parvocellular.

**Figure 4- supplement 2.**
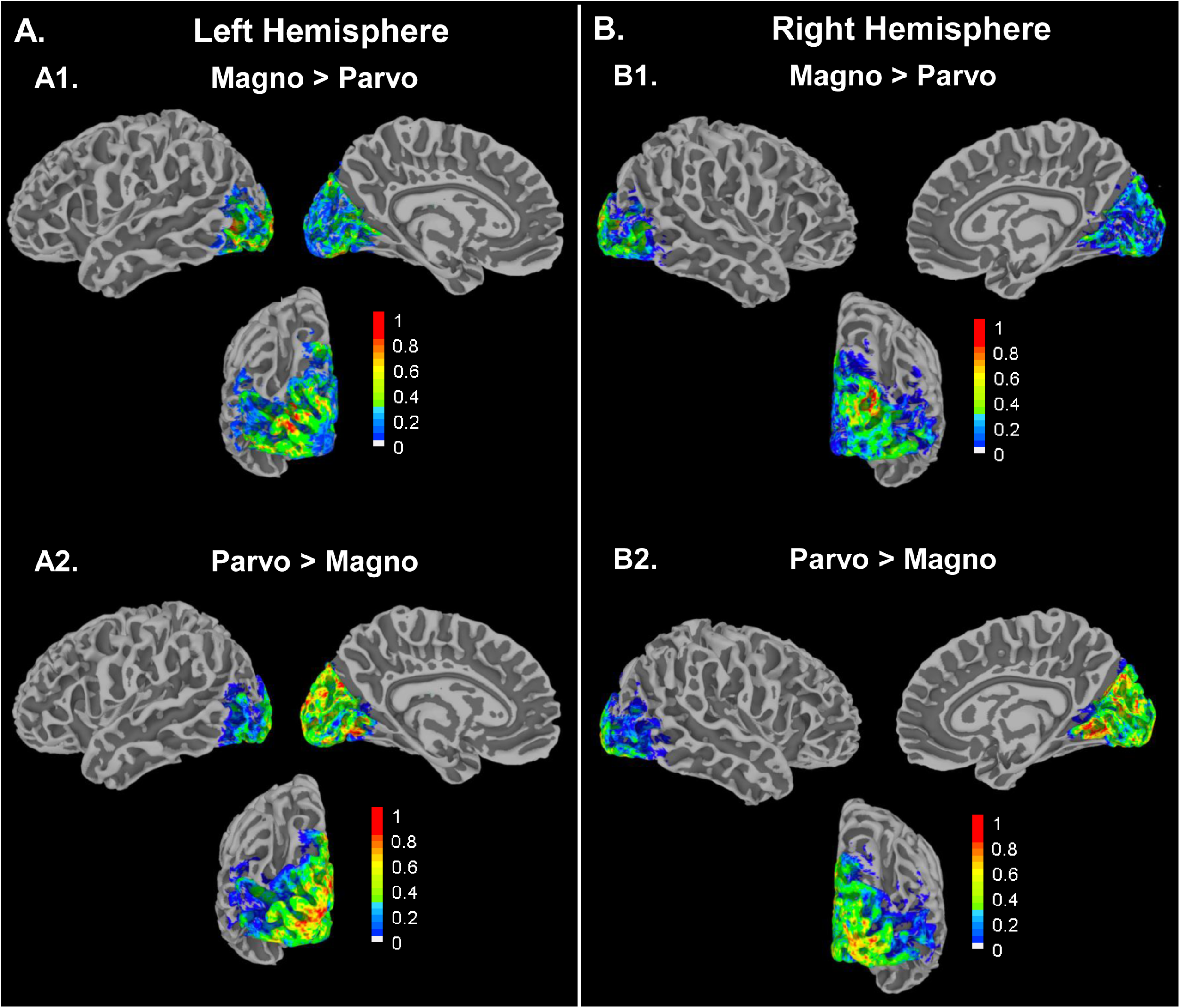
Retest session left and right hemisphere probabilistic maps showing the percentage of involvement of visual cortex voxels on surface renderings. A) Left hemisphere visual regions, with A1) showing the Magno > Parvo contrast and with A2) showing the Parvo > Magno contrast. B) Right hemisphere visual regions with B1) showing the Magno > Parvo contrast and with B2) showing the Parvo > Magno contrast. These probabilistic maps are not fully complementary in terms of voxel correspondence; while they represent opposite contrasts, they were generated by separated analyses. Magno = magnocellular; Parvo = parvocellular.

**Figure 5- supplement 1.**
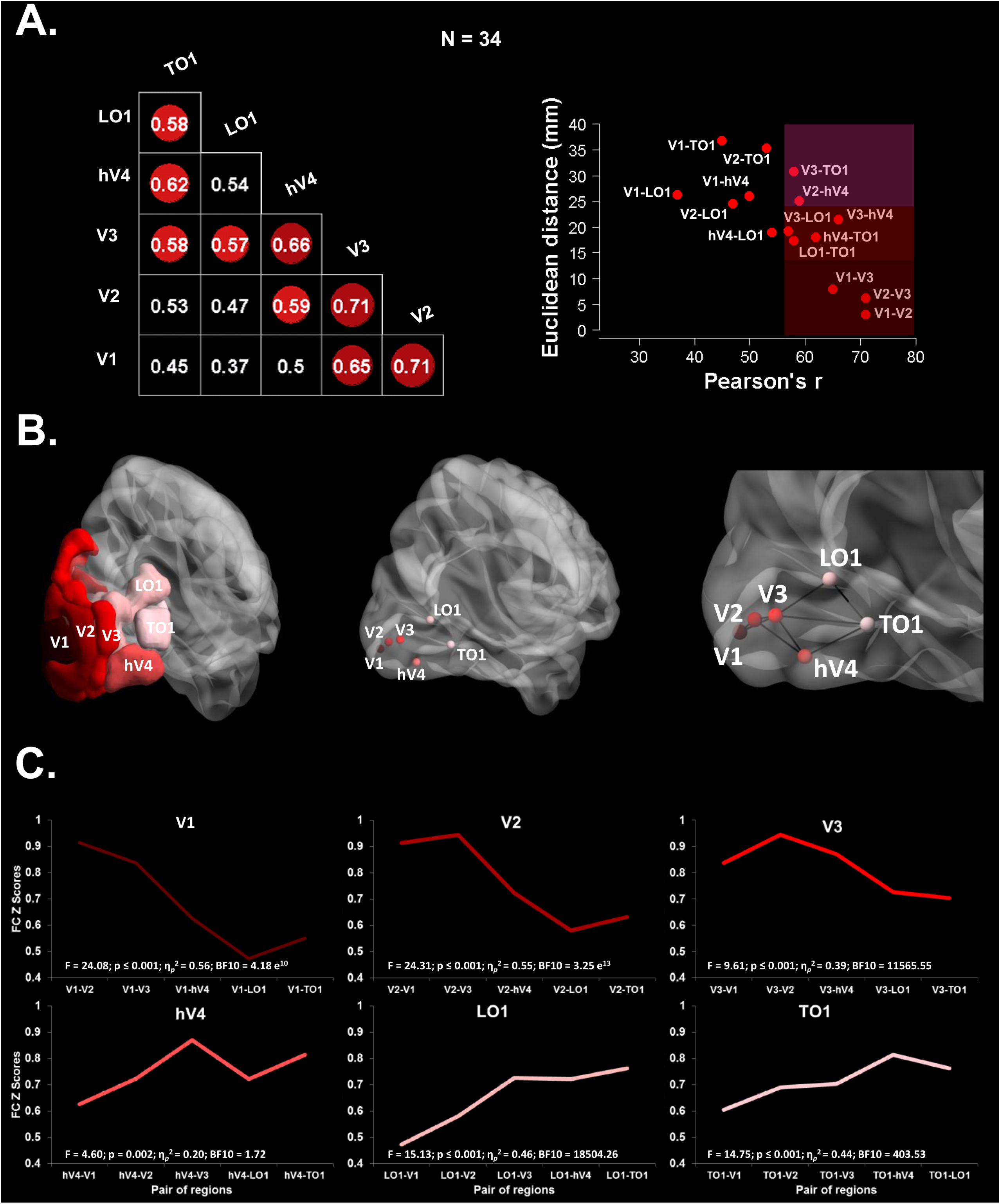
Right hemisphere task-related pairwise functional connectivity between visual cortex ROIs across experimental conditions (i.e., magnocellular-biased words, parvocellular-biased words, magnocellular-biased images, parvocellular-biased images. A) Left panel shows correlation matrix with significant Pearson’s r correlations (r ≥. 57) colored in red tones. Right panel shows Euclidean distances from posterior (V1, 0-value) to anterior (TO1) left visual cortex regions plotted against Pearson’s r values for each ROI pair, with shaded areas indicating pairs showing significant coupling. B) Left panel shows the sagittal rendering of the right visual cortex ROIs. Central panel shows the sagittal rendering of right visual cortex nodes centered at the ROIs center of mass. Right panel shows statistically significant pairwise correlations (i.e., edges) between right visual cortex nodes in a rendering of the posterior sagittal section. C) Task-related pairwise functional connectivity (Z scores) between right visual cortex ROI pairs across experimental conditions. The upper left panel shows Z scores between V1 and the rest of the ROIs, upper middle panel shows Z scores between V2 and the rest of the ROIs, and upper right panel shows Z scores between V3 an the rest of the ROIs. The lower left panel shows Z scores between hV4 and the rest of the ROIs, lower middle panel shows Z scores between LO1 and the rest of the ROIs, and lower right panel shows Z scores of TO1 and the rest of the ROIs.

**Figure 5- supplement 2.**
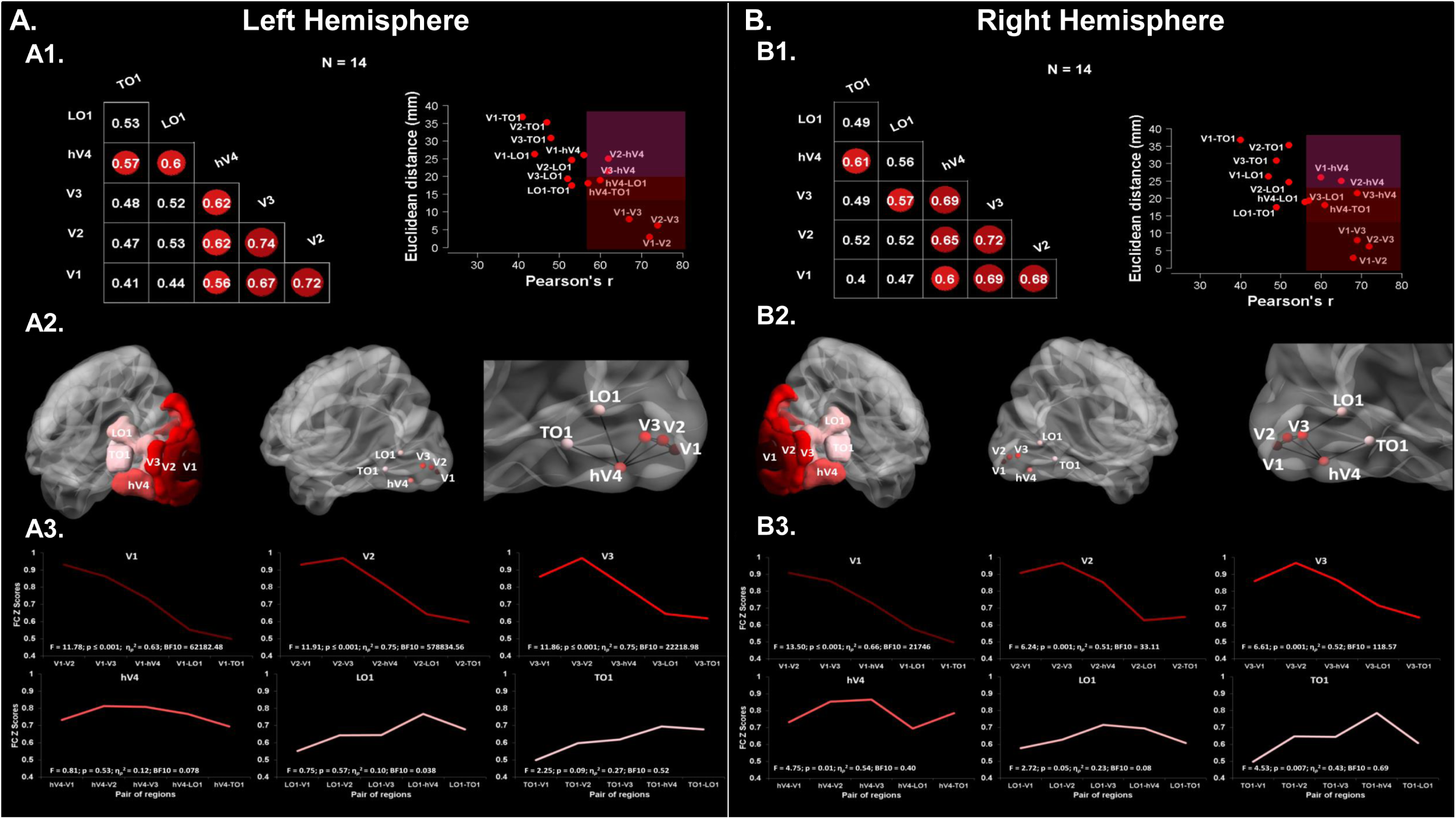
Retest session left and right hemisphere task-related pairwise functional connectivity between visual cortex ROIs across experimental conditions (i.e., magnocellular-biased words, parvocellular-biased words, magnocellular-biased images, parvocellular-biased images). A) Left hemisphere pairwise functional connectivity between visual cortex ROIs with A1) showing the correlation matrix with significant Pearson’s r correlations (r ≥. 57) colored in red tones (left panel) and Euclidean distances from posterior (V1, 0-value) to anterior (TO1) left visual cortex regions plotted against Pearson’s r values for each ROI pair, with shaded areas indicating pairs with significant coupling (right panel). A2) Left panel shows the sagittal rendering of the left visual cortex ROIs. Central panel shows the sagittal rendering of left visual cortex nodes centered at the ROIs center of mass. Right panel shows statistically significant pairwise correlations (i.e., edges) between left visual cortex nodes in a posterior sagittal rendering section. A3) Task-related pairwise functional connectivity (Z scores) between left visual cortex ROI pairs across experimental conditions. The upper left panel shows Z scores between V1 and the rest of the ROIs, upper middle panel shows Z scores between V2 and the rest of the ROIs, and upper right panel shows Z scores between V3 and the rest of the ROIs. The lower left panel shows Z scores of hV4 with the rest of the ROIs, lower middle panel shows Z scores between LO1 and the rest of the ROIs, and upper right panel shows Z scores between TO1 and the rest of the ROIs. A) Right hemisphere pairwise functional connectivity between right visual cortex ROIs with B1) showing the correlation matrix with significant Pearson’s r correlations (r ≥. 57) colored in red tones (left panel) and Euclidean distances from posterior (V1, 0-value) to anterior (TO1) right visual cortex regions plotted against Pearson’s r values for each ROI pair, with shaded areas indicating pairs with significant coupling (right panel). B2) Left panel shows the sagittal rendering of the right visual cortex ROIs. Central panel shows the sagittal rendering of right visual cortex nodes centered at the ROIs center of mass. Right panel shows statistically significant pairwise correlations (i.e., edges) between right visual cortex nodes in a posterior sagittal rendering section. B3) Task-related pairwise functional connectivity (Z scores) between right visual cortex ROI pairs across experimental conditions. The upper left panel shows Z scores between V1 and the rest of the ROIs, upper middle panel shows Z scores between V2 and the rest of the ROIs, and upper right panel shows Z scores between V3 and the rest of the ROIs. The lower left panel shows Z scores of hV4 with the rest of the ROIs, lower middle panel shows Z scores between LO1 and the rest of the ROIs, and upper right panel shows Z scores between TO1 and the rest of the ROIs.

